# seRNA *PAM-1* regulates skeletal muscle satellite cell activation and aging through *trans* regulation of *Timp2* expression synergistically with Ddx5

**DOI:** 10.1101/2021.10.06.463443

**Authors:** Karl Kam Hei So, Yile Huang, Suyang Zhang, Liangqiang He, Yuying Li, Xiaona Chen, Yu Zhao, Yingzhe Ding, Jiajian Zhou, Jie Yuan, Mai Har Sham, Hao Sun, Huating Wang

**Author notes:** These authors contribute equally to this paper as first authors (Karl Kam Hei So, Yile Huang). Co-correspondence, **Address correspondence to:** Huating Wang, 507A Li Ka Shing Institute of Health Sciences, Prince of Wales Hospital, The Chinese University of Hong Kong, Shatin, Hong Kong SAR, China., Hao Sun, 503, Li Ka Shing Institute of Health Sciences, Prince of Wales Hospital, The Chinese University of Hong Kong, Shatin, Hong Kong SAR, China.

## Abstract

Muscle satellite cells (SCs) are responsible for muscle homeostasis and regeneration; and lncRNAs play important roles in regulating SC activities. Here in this study, we identify *PAM-1* (Pax7 Associated Muscle lncRNA) that is induced in activated SCs to promote SC activation into myoblast cells upon injury. *PAM-1* is generated from a myoblast specific super-enhancer (SE); as a seRNA it binds with a number of target genomic loci predominantly in *trans*. Further studies demonstrate that it interacts with Ddx5 to tether *PAM-1* SE to it inter-chromosomal targets *Timp2 and Vim* to activate the gene expression. Lastly, we show that *PAM-1* expression is increased in aging SCs, which leads to enhanced inter-chromosomal interaction and target genes up-regulation. Altogether, our findings identify *PAM-1* as a previously unknown lncRNA that regulates both SC activation and aging through its *trans* gene regulatory activity.

## Introduction

Skeletal muscle tissue homeostasis and regeneration relies on muscle stem cells, also known as muscle satellite cells (SCs). These cells reside in a niche between the muscle fiber sarcolemma and the basal lamina surrounding the myofiber and are uniquely marked by transcription factor paired box 7 (Pax7). SCs normally lie in a quiescent state, upon activation by injury and disease, the cells quickly activate and express the master myogenic regulator, MyoD, then re-enter cell cycle and proliferate as myoblasts, subsequently differentiate and fuse to form myotubes (1, 2). A subset of SCs undergo self-renewal and return to quiescence, thus restoring the stem cell pool. Deregulated SC activity contributes to the development of many muscle associated diseases. For example, sarcopenia, a highly prevalent elderly disorder condition characterized by declined muscle mass and deficient muscle strength and function, is linked to a progressive reduction in the regenerative capacity of the SCs. It is thus imperative to understand the way SCs contribute to muscle regeneration, and their potential to cell-based therapies. At cellular level, every phase of SC activity is tightly orchestrated by many molecules and signalling pathways both intrinsically from the cell and extrinsically from the niche; the elucidation of factors and molecular regulatory mechanisms governing SC function thus is of extreme importance, being the first step toward successful use of these cells in therapeutic strategies for muscle diseases.

It has become increasingly clear that long non-coding RNAs (lncRNAs) are important players regulating SC regenerative activates (3). For example, *SAM* promotes myoblast proliferation through stabilizing Sugt1 to facilitate kinetochore assembly (4); *Linc-YY1* promotes myogenic differentiation and muscle regeneration through interaction with YY1 (5). In another example, we found that in myoblast cells master transcription factor MyoD induces the expression of lncRNAs from super enhancers (SEs), so called seRNAs, which in turn regulate target gene expression in *cis* through interacting with hnRNPL (6). In fact, the functional synergism between enhancer generated eRNAs and their associated enhancer activity in regulating target promoter expression is well established. There are myriad of mechanisms that eRNAs or lncRNAs cooperate with protein, DNA or RNA partners to regulate transcription of target genes either in *cis* and in *trans*. For example, it is known that specific eRNA can interact with CBP or BRD4 within topologically associating domain (TAD) in a localized manner (7–9). Similarly, eRNA *Sphk1* evicts CTCF, that insulates between enhancer and promoter, thus activating proto-oncogene *SPHK1* expression in *cis* (10). Some eRNAs or lncRNAs, on the other hand, play a dual molecular function both in *cis* and in *trans*, for example, *lincRNA-p21* acts in *trans* by recruiting heterogenous nuclear ribonucleoprotein K (hnRNPK) to the target promoter (11). Interestingly, in a separate study, *lincRNA-p21* can also transcriptionally activate *Cdkn1a* in *cis* (12). Another well demonstrated example of *trans* acting eRNA is *FIRRE*, that interacts with hnRNPU via a conserved RRD nuclear localization during hematopoiesis (13–15). Also, distal regulatory region of *MyoD* transcribed eRNA ^DRR^eRNA that interacts with cohesin and transcriptionally activates *Myogenin* in *trans* (16). Altogether, these findings demonstrate the diversified modes of action of eRNAs or lncRNAs in regulating target genes, which needs to be more exhaustively investigated.

Here in this study, we identify *PAM-1*, an seRNA that regulates SC activation. Expression of *PAM-1* is evidently upregulated during SC activation; consistently, knockdown of *PAM-1 in vitro* hindered SC activation. High throughput identification of *PAM-1* interactome reveals that it regulates Tissue inhibitor of metalloproteinases 2 (*Timp2*) locus in *trans* through binding and recruiting Ddx5 protein; loss of *PAM-1* or Ddx5 results in reduction of chromatin interaction between *PAM-1* SE and *Timp2* locus. Furthermore, *PAM-1* SE activity and chromatin connectivity with *Timp2* is elevated in aging SCs; *in vivo* inhibition of the SE activity by JQ1 reduces *Timp2* expression. Altogether our findings have identified *PAM-1* as a seRNA regulator of SC activation through its *trans* regulation of *Timp2* synergistically with Ddx5 to promote SC activation.

## Results

### lncRNAs profiling identifies *PAM-1* as a seRNA promoting SC activation

To gain global insights into the catalogue of lncRNAs in adult MuSC activation, we performed RNA-seq on quiescent satellite cells (QSCs) which were *in situ* fixed in ice-cold 0.5% paraformaldehyde before cell dissociation to preserve their quiescence, freshly isolated satellite cells (FISC), FISCs cultured and activated for 24 and 48 hours (ASC-24h and ASC-48h, respectively) (Fig 1A). lncRNAs expressed in each stage were identified and many lncRNAs were up- or down-regulated in ASC-24h or ASC-48h vs. QSCs (Fig 1B, and Suppl Table 1). 77 lncRNAs were up-regulated and 130 down-regulated in ASC-24h vs. FISC. (Suppl Figs 1A&B, Suppl Table 1). The above results suggested the dynamic expression of lncRNAs during SC activation. To further identify key lncRNAs that potentially play a role in SCs, we sought to identify Pax7 regulated lncRNAs by analyzing publicly available Pax7 ChIP-seq data in myoblast (17). Among 207 differentially expressed lncRNAs in ASC-24h vs FISC, we found that Pax7 binding was enriched in promoter regions of 32 of them (Suppl Table 1); we thus named them as Pax7 associated muscle lncRNAs (*PAMs*). Among them, *PAM-1* represented as one of the most up-regulated lncRNA in ASC-24h vs. FISC, with no expression in FISC but reaching 12.7 FPKM in ASC-24h (Suppl Figs 1B&C), suggesting it possibly promotes SC activation/proliferation. Previously known as *Gm12603, PAM-1* is a lncRNA located on chromosome 4: in the intervening region of Interferon alpha (*Ifna*) family and Cyclin-dependent kinase 2 inhibitor (*Cdkn2a* and *Cdkn2b*) protein coding genes. *Gm12603* is also known as *Wincr1*, a Wnt activated lncRNA in mouse dermal fibroblast affecting extracellular matrix composition via collagen accumulation in dermal fibrosis (18). To dissect its function in SCs, we next cloned its sequence from C2C12 myoblast cells by Rapid Amplification of cDNA ends (RACE); it was 718bp long with three exons. More interestingly, we found that *PAM-1* is generated from a SE region defined using our published H3K27ac ChIP-seq datasets (Fig 1C) (19). Concomitant with the expression pattern of *PAM-1*, high level of H3K27ac ChIP-seq signals were observed in ASC-24h but not in FISC (Fig. 1D). The above findings suggested that *PAM-1* may function as a seRNA to promote SC activation. Indeed, knockdown of *PAM-1* reduced the activation of SCs as revealed by the EdU assay (Fig. 1E). Overexpression of *PAM-1* in FISCs by transfection of a *PAM-1* expression plasmid resulted in 16.65% increase of Pax7 MyoD double positive ASCs, concomitant with reduction in number of Pax7+MyoD-QSCs by 16.65%, (Fig 1F). Similar phenomena were also observed in SCs associated with isolated single muscle fibers; overexpressing the *PAM-1* plasmid led to a 26.66% increase of Pax7+MyoD+ cells (Fig 1G). Taken together, these findings suggest *PAM-1* is a potential seRNA that promotes activation of SCs.

**Figure 1.**
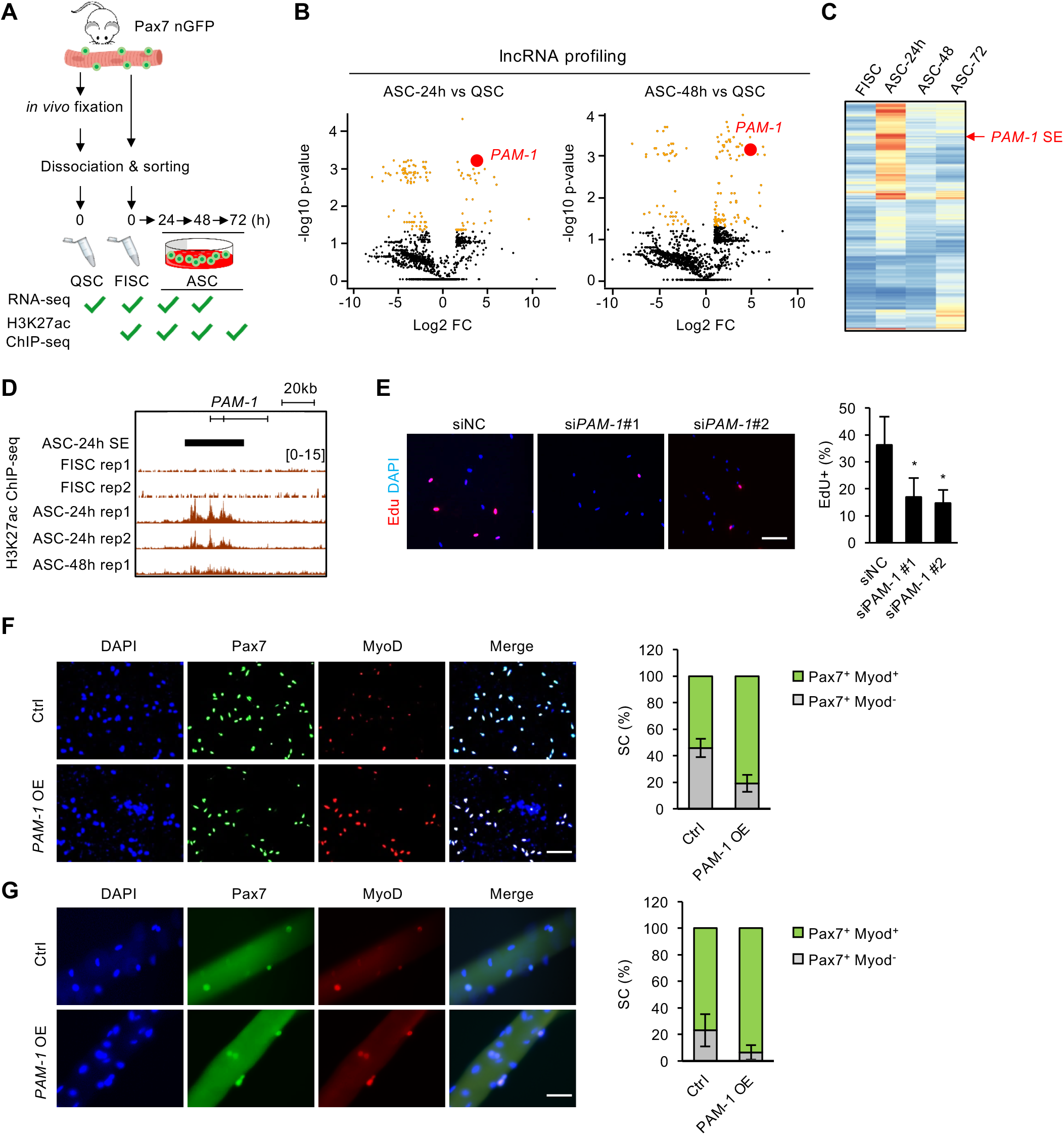
lncRNAs profiling identifies *PAM-1* as a seRNA promoting SC activation. **A.** Transcriptomic and epigenomic discovery of seRNAs in quiescence SCs (QSCs), freshly isolated SCs (FISCs), or activated SCs (ASCs) *in vitro* cultured for 24, 48 and 72 hours respectively. All SCs were isolated from muscles of Tg:Pax7-nGFP mice. **B.** Differentially expressed genes (DEGs) were identified in ASCs versus QSCs. Yellow dots indicate differentially expressed lncRNAs. *PAM-1* was highly expressed in ASCs. **C.** Heatmap showing H3K27ac signal intensity on super-enhancers along SC activation. *PAM-1* associated SE was inactive in FISCs, but the activity peaked in ASCs after 24 hours of *in vitro* culture, followed by a reduction in SE activity. **D.** Genome browser tracks showing active histone mark H3K27ac on *PAM-1* locus. **E.** Knockdown of *PAM-1* expression *in vitro* in cultured ASCs for 48 hours with *in vitro* EdU incorporation assay. The percentage of EdU+ cells was quantified. **F-G.** Overexpression of *PAM-1* in FISCs or freshly isolated myofibers increased the percentage of Pax7+Myod+ cells 48 hours post-transfection. The percentage of double positive cells was quantified. (Data represent the mean ± SD. P-value was calculated by two-tailed unpaired *t* test (*P < 0.05). Scale bars: 100μm.)

### *PAM-1* interacts with inter-chromosomal loci to modulate target expression

To further elucidate the regulatory mechanism of *PAM-1* in SC activation, we sought to identify the subcellular localization pattern of *PAM-1* as lncRNA function is largely determined by its cellular localization (20, 21) and seRNAs are known to be localized in both nucleus and cytoplasm of muscle cells (6). Cellular fractionation was performed using C2C12 myoblasts and *PAM-1* was found to localize largely in nuclear fraction (70.21%); as controls, lncRNAs *Xist* and *Malat1* were predominately nuclear localized (93.61% and 87.37% respectively) while *Gapdh* mRNAs were enriched in the cytoplasm (89.66%) (Fig 2A). The above finding was further validated by RNA Fluorescence in situ hybridization (FISH) using *PAM-1* anti-sense probe, which also revealed *PAM-1* transcript was predominantly enriched in the nucleus of ASCs (Fig 2B). Lastly, Subcellular localization of *PAM-1* was also confirmed using sucrose gradient centrifugation on C2C12 myoblast lysate which separated protein complex based on their size. Using cohesin loading factor NIPBL as positive control for nuclear fraction, we also found *PAM-1* mainly enriched in the nucleus of myoblasts (Fig 2C). Taken together, our results showed *PAM-1* is a nuclear enriched seRNA.

**Figure 2.**
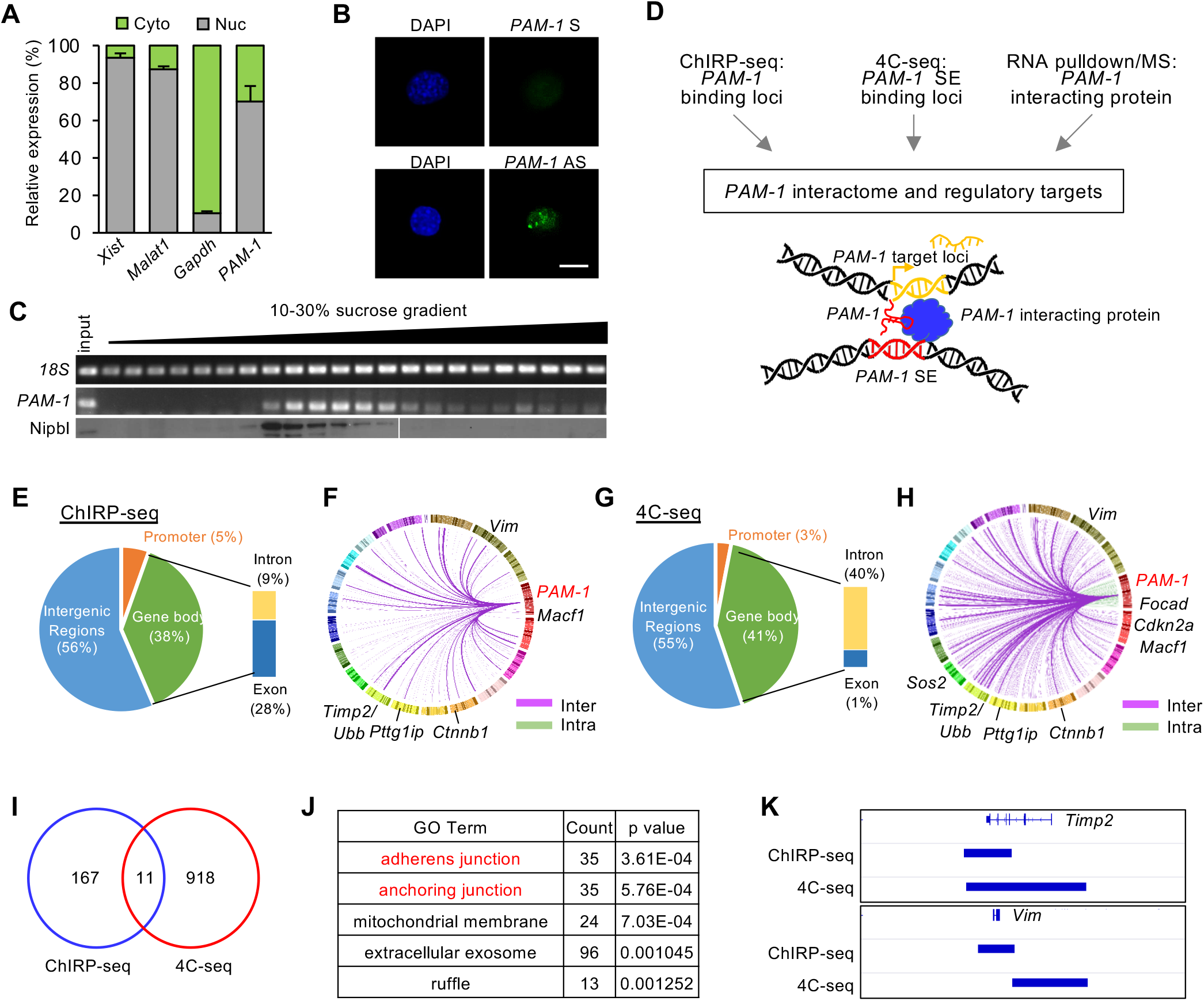
*PAM-1* is a nuclear-retained lncRNA, forming *cis* and *trans* chromosomal interactions. **A.** Cellular fractionation of C2C12 cell line showed *PAM-1* was enriched in nucleus not cytosolic fraction. *Xist* and *Malat1* were positive control for nuclear fraction, *Gapdh* was positive control for cytosolic fraction. **B.** Fluorescence in situ Hybridization (FISH) using *PAM-1* antisense (AS) probe showed nuclear localization of *PAM-1*, sense probe (S) was used as negative control. Scale bar: 10μm **C.** Cellular fractionation using sucrose gradient ultracentrifugation showed *PAM-1* was co-localized in fractions containing nuclear protein Nipbl. **D.** Experimental workflow to discover *PAM-1* interactome and regulatory targets. For details and controls, see Materials and Method. **E.** Pie chart showing distribution of *PAM-1* seRNA interacting chromatin across the genome in ChIRP-seq. **F.** Circos plot showing genes associated to *PAM-1* seRNA interacting chromatin, each line in the plot represents an interaction, line colors represent interchromosomal (purple) or intrachromosomal (green) interactions. Chromosome numbers were colored and arranged in clockwise direction. Top ranked genes were named in the figure. **G.** Pie chart showing distribution of *PAM-1* SE interacting chromatin across the genome in 4C-seq. **H.** Circos plot showing genes associated to *PAM-1* SE interacting chromatin, each line in the plot represents an interaction, line colors represent inter-chromosomal (purple) or intra-chromosomal (green) interactions. Chromosome numbers were colored and arranged in clockwise direction. Top ranked genes were named in the figure. **I.** Venn diagram showing overlapping loci between ChIRP-seq of *PAM-1* seRNA and 4C-seq of *PAM-1* SE. **J.** Table showing top ranked gene ontology (GO) terms of genes associated with overlapping loci from ChIRP-seq of *PAM-1* seRNA and 4C-seq of *PAM-1* SE. **K.** Genome browser tracks showing *Timp2* and *Vim* as examples of *PAM-1* inter-chromosomal targets.

To elucidate the functional mechanism of *PAM-1* as a nuclear seRNA, *PAM-1* seRNA and *PAM-1* SE interacting loci on the genome were identified respectively (Fig 2D). We first performed *PAM-1* Chromatin Isolation by RNA Purification (ChIRP-seq) in ASCs to identify its binding target loci. As a result, we found that *PAM-1* seRNA interacted with 178 DNA regions and associated with 1199 genes across the genome (Figs 2E&F, Suppl Fig 2A and Suppl Table 2). Strikingly, *PAM-1* dominantly associated with inter-but not intra-chromosomal regions (Fig 2F). Only 5 out of the 178 regions were found on chromosome 4 and not adjacent to *PAM-1* gene locus (Suppl Table 2). The above findings suggested *PAM-1* seRNA may predominantly target gene loci in *trans*. Gene Ontology (GO) analysis on the above identified *PAM-1* ChIRP-seq targets revealed that they were enriched for actin filament organization and regulation of cell shape (Suppl Fig 2A). Next, we set out to identify *PAM-1* SE target genes by performing circular chromosome conformation capture (4C-seq) using *PAM-1* SE region as a bait to query genome-wide chromatin interactions. Among 929 interacting targets, 74% (687) were in *trans* while only 26% (242) were in *cis* on chromosome 4 (Figs 2G&H and Suppl Table 3).

By comparing the above targets from *PAM-1* seRNA ChIRP-seq and *PAM-1* SE 4C-seq data, we identified 11 common loci in both datasets; one of the loci was located on chromosome 4, 33Mb downstream of *PAM-1* on chromosome 4, while others were all located in other chromosomes (Fig 2I), suggesting *PAM-1* and *PAM-1 SE* may regulate genes in *trans* together. These 11 loci were associated with 152 genes, which were enriched for GO terms such as “adherens junction”, “anchoring junction”, in which *Timp2* and Vimentin (*Vim*) genes were top ranked (Fig 2J&K). Timp2 plays a dual role in mediating extracellular matrix by mediating matrix metalloproteinase (MMP) activation and inhibition via interaction with MMP-14 and MMP-2 (22). Overexpression of *Timp2* in C2C12 myoblast was known to delay myogenic differentiation and arrest C2C12 in Myod+Myog-state (23). Vim is an intermediate filament protein to modulate cell shape and motility in myoblast, which is also considered as a reliable marker for regenerating muscle tissue (24–27). The above findings raised an intriguing possibility that *PAM-1* seRNA and *PAM-1* SE interact with the inter-chromosomal target loci *Timp2* and *Vim* to activate their expression. Consistent with the notion, we found that knocking down *PAM-1* seRNA using siRNA oligo decreased the expression levels of *Timp2* and *Vim* in C2C12 myoblasts (Fig 3A); removing *PAM-1* SE region using CRISPR/cas9 yielded similar molecular phenotype (Fig 3D). To test the possibility that *PAM-1* functions to tether *PAM-1* SE to the target loci, we performed 3C qRT-PCR assay in C2C12 myoblasts; in line with the 4C-seq result, and *PAM-1* locus indeed displayed evident interaction with *Timp2* and *Vim* promoters; however, *PAM-1* siRNA oligo mediated knockdown significantly reduced the interaction by 41.42% & 58.72% for *Timp2*, and 25.77% & 32.18% for *Vim* (Fig 3B). Consistently, down-regulation of *PAM-1* expression also led to a reduction in H3K27ac signals at *Timp2* and *Vim* loci (Fig 3C). Furthermore, knockout of the 59bp flanking 5’ of *PAM-1* transcript and the constituent enhancer of *PAM-1* SE region (Suppl Fig 2B) in C2C12 myoblast using CRISPR/Cas9 also led to reduction in the chromatin interaction between *PAM-1* SE with *Timp2* and *Vim*, and reduced expression of *Timp2* and *Vim* and associated H3K27ac signals (Figs 3C&F). Altogether, our results demonstrate that *PAM-1* seRNA and *PAM-1* SE can indeed interact with inter-chromosomal target loci *Timp2* and *Vim* to modulate their expression.

**Figure 3.**
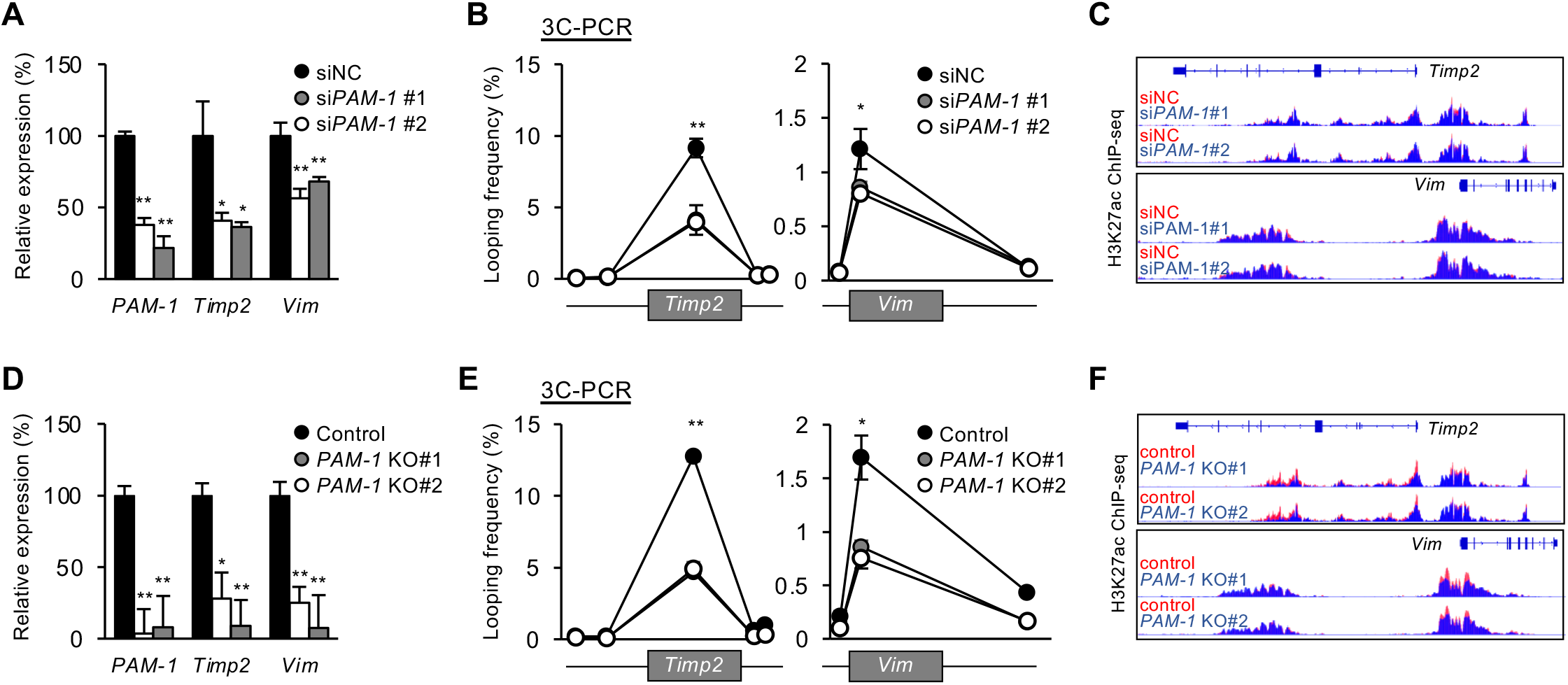
*PAM-1* regulates extracellular matrix associated genes *Timp2* and *Vim*. **A-C.** Knockdown of *PAM-1* expression with siRNA oligo treatment in C2C12 myoblast. **A.** Knockdown of *PAM-1* showed reduction in *Timp2* and *Vim* expression. **B.** Chromatin Conformation Capture assay (3C-qRT-PCR) with knockdown of *PAM-1* showed reduction in chromatin interaction between *PAM-1* SE with *Timp2* and *Vim*. **C.** Genome browser tracks showing knockdown of *PAM-1* led to mild reduction in H3K27ac signal intensity on *Timp2* and *Vim* loci. **D-F.** Knockout of *PAM-1* locus using CRISPR/cas9 approach in C2C12 myoblast. **D.** Knockout of *PAM-1* showed significant reduction in *Timp2* and *Vim* expression. **E.** 3C-qRT-PCR with knockout of *PAM-1* showed reduction in chromatin interaction between *PAM-1* SE with *Timp2* and *Vim*. **F.** Genome browser tracks showing knockout of *PAM-1* led to reduction in H3K27ac signal intensity on *Timp2* and *Vim* loci. Data information: Data represent the mean ± SD. P-value was calculated by two-tailed unpaired *t* test (*P < 0.05, **P < 0.01).

### *PAM-1* regulates inter-chromosomal targets via associating with Ddx5

To further explore molecular mechanism on how *PAM-1* regulates inter-chromosomal targets *Timp2* and *Vim*, we performed RNA-pulldown followed by mass spectrometry (MS) to identify its protein interactome (Fig 4A&B). Biotinylated sense probe of *PAM-1* was used in the RNA pull down *in vitro*, while biotinylated anti-sense probe was used as negative control for non-specific protein binding. Among 134 potential protein partners retrieved by the sense probe with at least 5 unique peptide count, two known RNA binding proteins (RBPs), DEAD-Box Helicase 5 (Ddx5) and DEAD-Box Helicase 17 (Ddx17) were highly ranked, which showed 22 and 14 unique peptide counts respectively with target protein size around 70kDa (Fig 4B). These proteins are commonly found to bind together in multiple cell types to carry out a myriad of molecular functions such as transcription regulation, rRNA processing, mRNA decay and splicing (28). To validate the above result, we performed RNA-pulldown followed by Western blot. *PAM-1* seRNA but not GFP control transcripts retrieved an evident amount of Ddx5 and Ddx17 from C2C12 myoblast (Fig 4C). To test if Ddx5 regulates target expression in cooperation with *PAM-1* seRNA, we first performed Ddx5 ChIP-seq which revealed an evident binding of Ddx5 on *Timp2* and *Vim* loci (Fig 4D), suggesting *PAM-1*/Ddx5 are both tethered to the target loci. Consistent with their possible functional synergism, knockdown of Ddx5 in ASCs led to down-regulation of *Timp2* and *Vim* (Fig 4E). Furthermore, knockdown of Ddx5 in ASCs reduced the interaction between *PAM-1* SE and promoters of *Timp2* and *Vim* (Fig 4F), suggesting Ddx5 promotes the inter-chromosomal interaction together with *PAM-1* (29). Lastly, knockdown of *Ddx5* reduced the number of activated SCs as revealed by the EdU assay (Fig 4G). Taken together, our findings demonstrate *PAM-1* seRNA and Ddx5 function synergistically in orchestrating the inter-chromosomal interactions between the *PAM-1* SE with the two target loci, *Timp2* and *Vim*, consequently promoting transcriptional activity.

**Figure 4.**
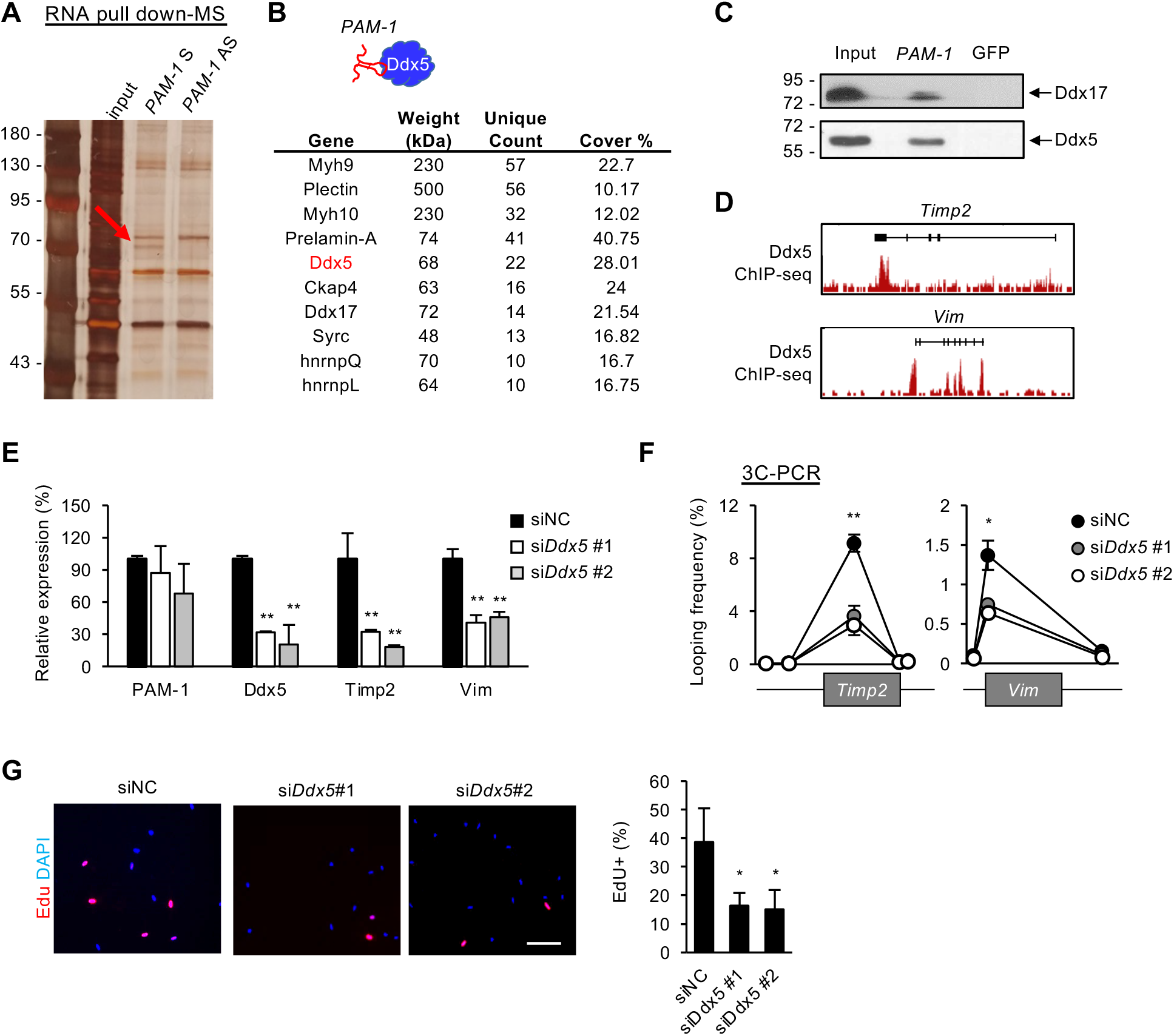
*PAM-1* regulates inter-chromosomal targets via association with Ddx5. **A.** RNA pulldown using *PAM-1* sense or antisense (AS) experiment followed by SDS-PAGE. AS was used as negative control. Red arrow indicates enrichment of a protein band at 70kDa specifically found in pulldown using *PAM-1* sense probe. **B.** Mass spectrometry (MS) result of the above band showing a list of potential protein binding partners of *PAM-1*. **C.** RNA pulldown followed by Western blotting of the above identified two candidate protein partners (Ddx5 and Ddx17) of *PAM-1* transcript. **D.** Ddx5 ChIP-seq in C2C12 myoblast showing enrichment of Ddx5 on promoter of *Timp2* or *Vim*. **E.** Knockdown of *Ddx5* using siRNA oligo in C2C12 myoblast showed reduction in expression of *Ddx5, Timp2* and *Vim* but not *PAM-1*. **F.** 3C-qRT-PCR with siRNA oligo mediated *Ddx5* knockdown showed reduction in chromatin interaction between *Timp2* or *Vim* promoter with *PAM-1* locus in the above cells. **G.** Knockdown of *Ddx5* expression *in vitro* in cultured ASCs for 48 hours showed reduction in EdU+ SCs. The percentage of EdU+ cells was quantified. Data information: Data represent the mean ± SD. P-value was calculated by two-tailed unpaired *t* test (*P < 0.05, **P <0.01). Scale bar: 100μm.

### *PAM-1* increase in aging SCs drives its target gene upregulation

Lastly, to test the possible involvement of *PAM-1* in SC aging, we performed ChIP-qPCR in ASCs from mice of various ages (2, 16 or 24 months) and found the activity of *PAM-1* SE was indeed increased by 13.63% at 16 months, and reached a plateau (26.45%) at 20 months (Fig 5A). The expression of *Timp2* also showed an increase (781.36%) in ASCs from 20 vs 2-month-old mice; but *Vim* expression remained unchanged (Fig 5B). Furthermore, the interaction between *PAM-1* locus and promoters of *Timp2* and *Vim* were found to increase by 46.75% and 50.18% respectively in 20 vs 2-month-old ASC (Fig 5C). Altogether, the above data suggest that *PAM-1* SE activity is elevated in aging SCs and it is accompanied by the enhanced inter-chromosomal interaction between *PAM-1* and target loci and upregulated target expression. Our result suggested *Timp2* chromatin activity was increased during aging, and potentially mediated by *PAM-1*. Lastly, to further confirm the synergistic function of *PAM-1* and Ddx5 in increasing *Timp2* expression in aging SCs, we found that knockdown of *PAM-1* or *Ddx5* in ASC in 20- or 30-month but not 2-month-old mice led to down-regulation of *Timp2* in ASCs (Fig 5D). Recently we have reported the use of BET family of bromodomain protein binding inhibitor JQ1 to down-regulate enhancer activity in aging mouse muscle (30). Expectedly, *in vivo* JQ1 treatment in 10-month-old mice led to a down-regulation of *PAM-1* seRNA and *Timp2*, but interestingly not *Vim* (Fig 5E), reinforcing the notion that *PAM-1* SE activation causes *Timp2* upregulation during SC aging.

**Figure 5.**
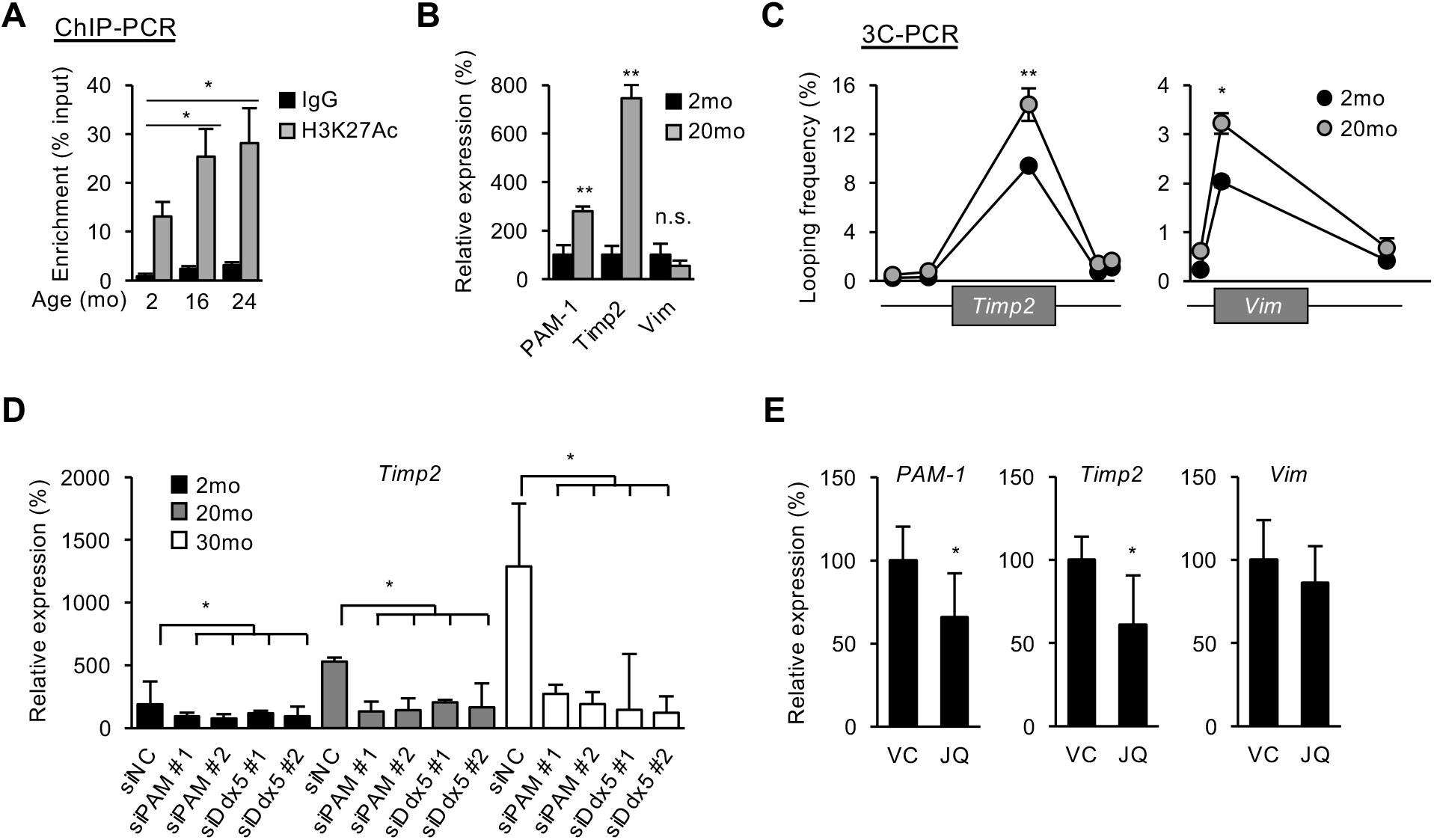
*PAM-1* increase in aging SCs drives its target gene upregulation. **A.** H3K27ac ChIP qRT-PCR showing increase in enrichment on *PAM-1* SE in ASCs isolated from aging (16 and 24 months) vs young (2 months) old mice. **B.** qRT-PCR showed up-regulation of *PAM-1* and target genes *Timp2* but not *Vim* in ASCs from 20 vs. 2 month old mice. **C.** 3C-qRT-PCR assay showed increase in interaction between *PAM-1* locus and *Timp2* or *Vim* promoter in the above aging ASCs. **D.** qRT-PCR showing the expression dynamics of *Timp2* in ASCs from 2, 20 and 30 months old mice. Knockdown of *PAM-1* or *Ddx5* using siRNA oligo in ASCs from 20 or 30 months old mice showed down-regulation of *Timp2*. **E.***In vivo* treatment of JQ1, a Brd4 inhibitor, in 10 month old mice down-regulated expression of *PAM-1* and *Timp2* but not *Vim* in FISCs. Data information: Data represent the mean ± SD. P-value was calculated by two-tailed unpaired *t* test (*P < 0.05, **P < 0.01).

## Discussion

Myogenesis is a complex process that relies on tightly regulated and finely tuned transcriptional regulatory mechanisms. Previous studies have discovered a myriad of lncRNAs that are dynamically regulated during myogenesis (31). Yet, their functional mechanisms in SCs remain largely unexplored. Here in this study we identified *PAM-1*, a seRNA that binds with Ddx5 protein to synergistically tether the SE to its inter-chromosomal target loci *Timp2* and *Vim*. Furthermore, we showed that deregulation of *PAM-1* in aging SCs drives the target gene deregulation (Fig 6).

**Figure 6.**
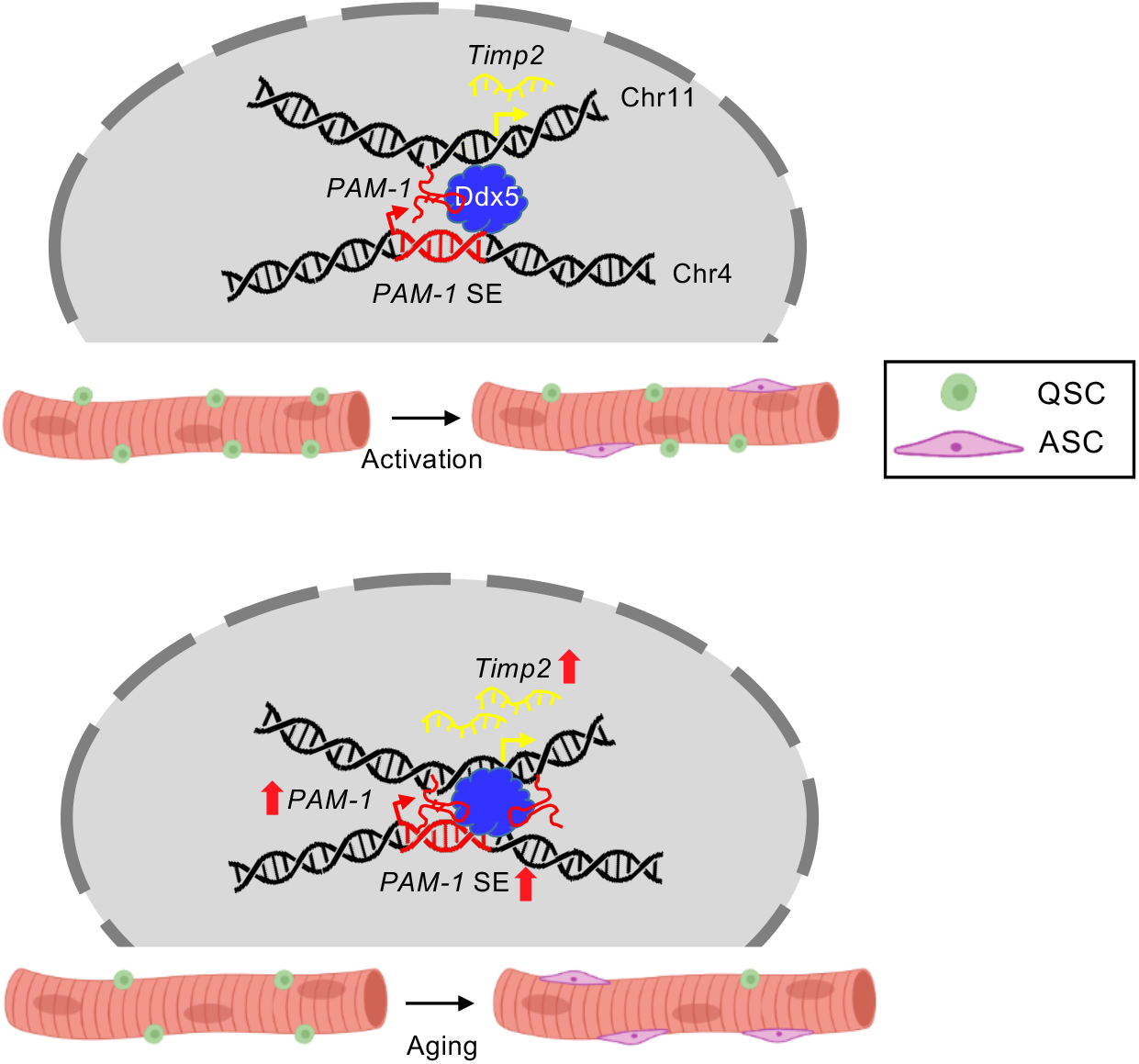
Schematic model showing functional role of seRNA *PAM-1* in SC activation and skeletal muscle aging. *PAM-1* regulates SCs activation by binding with Ddx5 to facilitate the chromatin interaction between *PAM-1* SE and target loci, *Timp2* and *Vim* during SC activation. In aging mice, the activity of *PAM-1* SE was elevated, thereby enhancing the transcription of *Timp2*, which potentially modulates extracellular matrix components in skeletal muscle.

Through transcriptomic profiling, *PAM-1* was identified as highly induced in activated SCs. Loss- and gain-of-function studies in SCs indeed pinpointed it as a promoting factor for SC activation. Its nuclear localization is consistent with its nature of being a seRNA. As part of the SE regulatory machinery, *PAM-1* and its residing SE together promote the expression of their target loci, *Timp2* and *Vim*, both encoding extracellular matrix proteins. Increased extracellular matrix (ECM) proteins expressions are essential to SC activation (32). The regulation of ECM composition was conventionally believed to be mediated by surrounding fibroblasts, fibro/adipogenic progenitors, myofibers and basal lamina. In addition to receiving signals from ECM, emerging evidence demonstrates SCs also contribute to ECM compositions through secretion of matrix metalloproteases and urokinase plasminogen activator (33, 34). For example, transcriptome profiling of freshly isolated SC revealed that cell adhesion and ECM genes, such as Timp and integrin, were differentially expressed when compared with freshly isolated SC from dystrophic mdx mice (35, 36). This correlation was further demonstrated by systemic delivery of MMP inhibitor, AM409, which impairs SC activation (35). Therefore, upregulation of ECM gene expression mediates the promoting function of *PAM-1* during SC activation. Furthermore, we showed that *PAM-1* upregulation contributes to ECM increase in aging SCs. It is known that the regenerative potential of SCs declines during aging, which is in concurrent with its fibrogenic conversion and muscle fibrosis (37). Our findings thus provide a potential way to partially restore ECM in aging SCs by down-regulating *PAM-1* expression. Recently, an integrated transcriptome analysis of aging human skeletal muscle revealed a group of differentially expressed lncRNAs, and overexpression of lncRNA *PRKG1-AS1* could increases cell viability and reduces apoptosis in human skeletal myoblast (38). In the future more efforts will be needed to elucidate the potential roles of lncRNAs in aging SCs.

Mechanistically, our data highlights the important role of *PAM-1* seRNA to regulate inter-chromosomal targets through tethering its residing SE to the target loci. eRNAs or seRNAs are commonly known to regulate enhancer-promoter interactions as an integrated component of SE activating machinery (6); it is believed that *cis* regulation of the target loci through intra-chromosomal interactions is a more prevalent mode compared to *trans* regulation via inter-chromosomal interactions (39). However, *trans* activating eRNAs do exist to translocate to distal chromosomal regions beyond its neighboring loci. For example, *MyoD* distal enhancer generates ^DRR^eRNA that transcriptionally regulate *Myogenin* expression in *trans* via cohesin recruitment (16). Similarly, in human prostate cancer, an adjacent eRNA of kallikrein related peptidase 3 (*KLK3*) can regulate target genes expression in *trans* (40). Our findings from integrating ChIRP-seq and 4C-seq demonstrate that *PAM-1* mainly acts in *trans* to exert its regulatory function in SCs; thus provide additional evidence to support the *trans* regulatory mechanism by eRNAs.

Another important discovery from our study stems from the identification of a direct physical interaction between *PAM-1* and Ddx5. Ddx5 and *PAM-1* synergistically facilitate the SE-target interaction; knockdown of *Ddx5* impaired the interaction. Although classically known as a RNA helicase controlling mRNA splicing, recent studies demonstrated Ddx5 interacts with a myriad of lncRNAs, and uses the lncRNAs as a scaffold to bring in specific transcriptional machinery or chromatin architectural protein in context dependent manner (28). It was also demonstrated that Ddx5 interacts with lncRNA *mrhl* to mediate cell proliferation in mouse spermatogonial cells (41). Ddx5 and Ddx17 (p68 and p72) bind with lncRNA *SRA* in regulating skeletal muscle differentiation (42). With foundation laid by these studies, we further demonstrated the functional role of Ddx5 to mediate chromatin interactions via its interaction with *PAM-1*, underscoring the prevalence of lncRNAs and RNA helicases interaction and also broadening the mechanisms through which lncRNAs regulate gene expression.

## Materials and Method

### Mice

Tg:Pax7-nGFP mouse strain (43) was kindly provided by Dr. Shahragim Tajbakhsh. All animal handling procedures and protocols were approved by the Animal Experimentation Ethics Committee (AEEC) at the Chinese University of Hong Kong.

### JQ1 treatment

JQ1 treatment was performed as described previously (30). C57/BL6 mice were caged in groups of five and maintained at controlled temperature (20±1°), humidity (55±10%), and illumination with 12hours light/ 12hours dark cycle. Food and water were provided *ad libitum*. All procedures involving animal care or treatments were approved by the Animal Ethics Committee (AEEC) at Chinese University of Hong Kong (CUHK) on the protection of animals used for scientific purposes. To investigate the effect of JQ1 on aging skeletal muscle, daily intraperitoneal injection of JQ1 at 50mg/kg was performed on 10-month-old mouse for 14 days (30), with DMSO as control. Then TA muscle tissues were extracted from mice, and MuSCs were isolated from mouse skeletal muscle by fluorescence-activated cell sorting (FACS) using BD FACSAria Fusion cell sorter (BD Biosciences) with cell surface marker Sca1^-^/CD31^-^/CD45^-^/Vcam^+^ (44).

### Cells

Mouse C2C12 myoblast cell (CRL-1772) was obtained from American Type Culture Collection (ATCC) and cultured in DMEM medium with 10% fetal bovine serum, 1% penicillin/ streptomycin at 37°C in 5% CO_2_. Oligonucleoties of siRNA against mouse *PAM-1* and scrambled control were obtained from Ribobio Technologies (Guangzhou, China). siRNAs were transfected at 100nM into C2C12 using Lipofectamine 2000 (Life Technologies). The sequences of oligonucleotides using for siRNA knockdown were listed in Supplementary Table 4.

### Satellite cell isolation and culture

Hindlimb skeletal muscles from Tg:Pax7-nGFP mice were dissected and minced, followed by digestion with Collagenase II (LS004177, Worthington, 1000 units/mL) for 90 min at 37°C in water bath shaker. Digested muscles were then washed in washing medium (Ham’s F-10 medium (N6635, Sigma) containing 10% heat inactivated horse serum (Gibco, 26050088) with 1% penicillin/ streptomycin, followed by incubating in digestion medium with Collagenase II (100 units/mL) and Dispase (1.1 unit/mL, Gibco, 17105-041) for additional 30 min. Suspensions were then passed through 20G syringe needle to release myofiber-associated SCs. Mononuclear cells were filtered with a 40μm cell strainer, followed by cell sorting using BD FACSAria Fusion Cell Sorter (BD Biosciences). BD FACSDiva (BD Biosciences, version 8.0.1) software was used to manage machine startup, data acquisition and analysis of flow cytometry data. Culture dish were coated with poly-D-lysine (Sigma, P0899) and Matrigel (BD Bioscience, 356234). FACS isolated SCs were seeded in coated culture dish and cultured in Ham’s F10 medium with 10% heat inactivated horse serum, 5ng/mL FGF-Basic (AA 10-155) (Gibco, PHG0026), or cultured in differentiation medium (Ham’s F10 medium with 2% horse serum and 1% penicillin/ streptomycin).

### EdU incorporation assay

EdU incorporation assay was performed as described previously (4). EdU was added to cultured MuSCs for 4 hours, followed by fixation in 4% paraformaldehyde (PFA) for 15 min and stained according to the EdU staining protocols provided by manufacturer (Thermo Fisher Scientific, C10086)

### Single myofibers isolation and culture

Externsor digitorum longus (EDL) muscles were dissected and digested in collagenase II (800 units/mL) in DMEM medium at 37°C for 75 min. Single myofibers were released by gentle trituration with Ham’s F-10 medium with heat inactivated horse serum and 1% penicillin/ streptomycin, then cultured in this medium for the follow up experiments.

### Genomic editing by CRISPR-Cas9 in C2C12 cells

To delete *PAM-1* exon1, target-specific guide RNAs (gRNAs) were designed using CRISPR design tool (http://crispr.mit.edu), followed by cloning into BbsI digested px330 plasmid (Addgene, 42230). To perform genomic deletion, a pair of gRNAs containing plasmids were co-transfected into C2C12 cells with screening plasmid pSIREN-RetroQ (Clontech) using Lipotectamine 2000. Cells were selected with 2.5μg of puromycin for 3 days at 48 hours post-transfection. Cells were diluted to 1 cell per well in 96 well plate. Individual colonies were PCR validated. Sequences of gRNAs and genotyping primers were listed in Supplementary Table 4.

### Plasmids

Full length cDNA of *Gm12603* (*PAM-1*) was cloned into pcDNA3.1 vector using HindIII and KpnI restriction enzymes digestion site. Primer sequences were listed in Supplementary Table 4.

### qRT-PCR

RNAs were extracted using Trizol (Life Technologies), followed by reverse transcription using SuperScript III Reverse Transcriptase (Life Technologies). PCRs were performed with SYBR green (Life Technologies) using 1μL of immunoprecipitated DNA as template. PCR products were analyzed by LC480 II system (Roche). Primers used are listed in Supplementary Table 4.

### RNA pulldown

RNA pulldown was performed as described previously (4). *PAM-1* DNA constructs were first linearized by single restriction enzyme digestion (NotI and XhoI for antisense and sense transcription respectively). Biotinylated transcripts were generated by these digested constructed by *in vitro* transcription using Biotin RNA labeling Mix (Roche) and MAXIscript T7/T3 *In Vitro* Transcription Kit (Ambion). Transcribed RNAs were denatured at 90°C for 2 minutes, then cooling on ice for 5 minutes, followed by addition of RNA structure buffer (Ambion) and refolding at room temperature for 20 minutes. Nuclear proteins from C2C12 cells were collected by resuspending cell pellet in nuclear isolation buffer (40mM Tris-HCl pH 7.5, 1.28M sucrose, 20mM MgCl_2_, 4% Triton X-100 and 1x protease inhibitor). Nuclei were collected by centrifugation at 3,000g and 4°C for 10 minutes. Supernatant was removed, and nuclear pellet was resuspended in 1mL RIP buffer (25mM Tris-HCl pH 7.4, 150mM KCl, 0.5mM DTT, 0.5% NP-40, 1mM PMSF, 1x RNase inhibitor and 1x protease inhibitor), followed by homogenization for 10 cycles (15seconds on/off) using Ika homogenizer (Ika-Werk Instruments, Cincinnati). Nuclear envelops and debris were removed by centrifugation at 16,200g for 10 minutes. For RNA pulldown assay, 1mg of nuclear extracts were incubated with 3μg of refolded RNA on rotator at room temperature for 1 hour. 30μL of prewashed Dynabeads M-280 Streptavidin were added to each reaction with incubate on rotator at room temperature for additional 1 hour. Streptavidin beads were collected using a magnetic rack, and beads were washed with 1mL RIP buffer for 5 times. Proteins were eluted by adding Western blot loading buffer and incubated at 95°C for 5 minutes, followed by removal of beads using magnetic rack. RNA pulldown samples were analyzed by SDS-PAGE followed by silver staining and LC-MS/MS with Q Exactive and Easy-nLC 1000 system (Thermo Fisher). Peptides were identified using MASCOT.

### RNA Fluorescence in situ hybridization (FISH)

RNA FISH was performed as described previously (45). Cells were fixed with 4% formaldehyde in PBS for 15 minutes at room temperature, followed by permeabilization with 0.5% Triton X-100, 2mM VRC (NEB) on ice, and two times 2x SSC wash for 10 minutes each. Probes were first amplified with PCR using *PAM-1* expression plasmid in RNA pulldown experiment. PCR products were then precipitated by ethanol, nick-translated and labelled with Green d-UTP (Abbott) and nick translation kit (Abbott). For each FISH experiment, 200μg of probe and 20μg of yeast tRNA were lyophilized and redissolved in 10μL formamide (Ambion), followed by denaturation at 100°C for 10 minutes and chilled immediately on ice. Denatured probes were mixed with hybridization buffer at 1:1 ratio. 20 μL of hybridization mix was added onto fixed cells, followed by putting coverslip on it and incubated at 37°C for 16 hours in a humidified chamber. Cells were then washed twice in 2x SSC, 50% formamide; thrice in 2x SSC; and once in 1x SSC for 5 minutes each in 42°C. Cells were mounted by coverslip with ProLong Gold Antifade Reagent with DAPI (Invitrogen). Fluorescence images were taken in Olympus microscope FV10000 and FV10-ASW software (version 01.07.02.02, Olympus).

### Cellular fractionation

Cellular fractionation was performed as described previously (45). C2C12 cell pellet from 1×10^6^ cells was lysed with lysis buffer (140mM NaCl, 50mM Tris-HCl pH 8.0, 1.5mM MgCl_2_, 0.5% NP-40, and 2mM Vanadyl Ribonucleoside Complex) for 5 minutes at 4°C, followed by centrifugation at 4°C 300g for 2 minutes. The supernatant after centrifugation was considered as cytoplasmic fraction and stored in −20°C for storage, while the pellet was resuspended in 175μL resuspension buffer (500mM NaCl, 50mM Tris HCl pH 8.0, 1.5mM MgCl_2_, 0.5% NP-40, 2mM Vanadyl Ribonucleoside Complex) and incubate at 4°C for 5 minutes. Nuclear-insoluble fraction in the resuspended pellet was removed by centrifugation at 4°C and 16000g for 2 minutes. RNA was extracted from cytoplasmic and nuclear soluble fraction by Trizol (Life Technologies).

### Sucrose gradient

Sucrose gradient was performed as described previously (46). C2C12 cells were lysed in cell lysis buffer (50mM Tris-HCl pH 7.6, 1mM EDTA, 1% Triton X-100, 10% glycerol, 1mM DTT, 1mM PMSF, 1x RNase inhibitor and 1x protease inhibitor). 500μL of whole cell lysate was added to 13.5mL of 10-30% sucrose gradient, followed by centrifugation at 38000 RPM at 4°C for 16 hours. The centrifuged lysate was fractionated in 500μL portion. To avoid cross-contamination, only odd-numbered fractions were obtained. Protein samples were resolved in SDS-PAGE, followed by western blotting. RNA samples were subjected to RT-PCR of *PAM-1* and *18S*, followed by 2% agarose gel electrophoresis.

### Chromatin Immunoprecipitation using sequencing (ChIP-seq)

ChIP assays were performed as previously described (47). C2C12 cells were crosslinked with 1% formaldehyde at room temperature for 10 minutes, followed by quenching with 0.125M glycine for 10 minutes. Chromatin was fragmented using S220 sonicator (Covaris), followed by incubation with 5μg of antibodies and 50μL Dynabeads Protein G magnetic beads (Life Technologies) at 4°C on rotator for overnight. Anti-histone H3-K27 acetylation (Abcam, ab4729), anti Ddx5 (Abcam, ab21696) and normal rabbit IgG (Santa Cruz Biotechnology, sc-2027) were used in ChIP assay. Beads were washed with 1mL RIPA buffer for 5 times, followed by decrosslinking at 65°C for 16 hours and DNA extraction with phenol/chloroform. Immunoprecipitated DNA was resuspended in 50μL of water. 200ng of immunoprecipitated DNA was used as starting material for NEBNext®Ultra II DNA Library Preparation kit for Illumina (NEB) according to manufacturer’s guideline. DNA libraries were sequenced in Illumina NextSeq 550 platform.

### Chromatin Isolation by RNA Purification using sequencing (ChIRP-seq)

Biotin labelled probes targeting *PAM-1* lncRNA were designed by ChIRP Designer (LGC Biosearch Technologies) and listed in Supplementary Table 4. Cells were rinsed in PBS, trypsinized, washed once with complete DMEM followed by resuspension in PBS. 10 million of ASCs were collected per ChIRP experiment for separated odd and even probe pools. Cell pellets were cross-linked with 1% Glutaraldehyde in 40mL PBS on rotator for 10 minutes at room temperature, followed by quenching the cross-linking reaction with 2mL 1.25M Glycine for 5 minutes and resuspend in 1mL chilled PBS. Cell pellets were collected at 2000RCF for 5 minutes at 4 °C, followed by removing PBS, snap frozen with liquid nitrogen and stored at −80°C. Cell pellets were lysed and sonicated according to our standard ChIP-seq protocol (47), then aliquoted into two 1mL samples. Before ChIRP experiment, DNA were extracted for quality control with size ranging from 100-500bp. For ChIRP experiment, 10μL of lysate were saved for DNA input. 1mL of sonicated lysate was mixed with 2mL of hybridization buffer (750mM NaCl, 50mM Tris-HCl pH7, 1mM EDTA, 1% SDS, 15% Formamide, 1x protease inhibitor and 1x RNase inhibitor). 100pmol of odd and even ChIRP probes were added separately to the hybridization mixture and incubate at 37°C for 4 hours with rotation. After the hybridization was completed, 100μL of streptavidin magnetic C1 beads (Life Technologies, 65001) were washed thrice with hybridization buffer and added to each ChIRP reaction for extra 30 minutes incubation at 37°C with rotation. After the hybridization completed, 1mL of wash buffer (2X SSC, 0.5% SDS and 1x protase inhibitor) was used to wash the beads for 5 times using magnetic stand. Input control and *PAM-1* bound DNA was eluted with ChIP elution buffer for each pair of ChIRP reactions using standard elution protocol as ChIP (47). For ChIRP-seq, DNA libraries were prepared as previous described in ChIP-seq protocol (47). Raw reads were uniquely mapped to mm9 reference genome using Bowtie2 (48). Peaks were called by using MACS2 (49).

### 4C-seq and 3C qRT-PCR

3C experiments were performed as previously described using restriction enzyme Bgl II to digest fixed chromatins (46). Primers for *PAM-1* bait region and target regions were listed in Supplementary Table 4. First round of restriction enzyme digestion in 4C-seq was the same as 3C qRT-PCR. 4C experiment was then continued with TatI restriction enzyme digestion, incubated overnight at 37°C and circularized using T4 DNA ligase. Gradient range of annealing temperature (55-65°C) were used to determine the optimum annealing temperature for inverse PCR. Primer sequences for inverse PCR were listed in Supplementary Table 4. PCR products were subject to standard sequencing library preparation as ChIP-seq and ChIRP-seq. Sequencing reads with 5’end matching the forward inverse PCR primer sequence were selected and trimmed, remaining sequences containing TatI sites were mapped to mm9 assembly using Bowtie2 (48) and the interaction regions are identified by fourSig (50).

### RNA-seq

Total RNAs were extracted using Trizol, followed by poly(A) selection (Ambion, 61006) and library preparation using NEBNext Ultra II RNA Library Preparation Kit (NEB). Barcoded libraries were pooled at 10pM and sequenced on Illumina HiSeq 1500 platform.

### Statistical analysis

Statistical analysis of experimental data was calculated by the Student’s t-test, whereas * P<0.05, ** P<0.01, and n.s. means not significant (P>=0.05).

### Data availability

RNA-seq, H3K27ac ChIP-seq, *PAM-1* ChIRP-seq and *PAM-1* 4C-seq using in this study have been deposited in Gene Expression Omnibus (GEO) database under the accession code (GSE180073).

## Author contributions

K.K.H.S., H.S. and H.W. designed the experiments; K.K.H.S., Y.H. and S.Z. conducted the experiments; L.H. provided support on CRISPR/cas9 experiments; Y.L. provided support on RNA pulldown experiments; X.C. provided support on cellular fractionation and RNA FISH; Y.Z. provided support on ChIRP-seq experiments; Y.D., J.Z. and J.Y. provided support on bioinformatics analysis; Y.H. analyzed the sequencing data; S.Z. contributed to *ex vivo* muscle fiber culture, M.H.S. provided resources for molecular experiments, K.K.H.S. and H.W. wrote the paper.

## Acknowledgements

We thank Prof. Zhenguo Wu, Prof. Danny CY Leung and Prof. Tom HT Cheung for their kind suggestions on skeletal muscle satellite cells and epigenomics. This work was supported by General Research Funds (GRF) from the Research Grants Council (RGC) of the Hong Kong Special Administrative Region (14116918, 14120420, and 14120619 to H.S.; 14115319, 14100018, 14100620, 14106117 and 14106521 to H.W.); the National Natural Science Foundation of China (NSFC) to H.W. (Project code: 31871304); Collaborative Research Fund (CRF) from RGC to H.W. (C6018-19GF); NSFC/RGC Joint Research Scheme to H.S. (Project code: N_CUHK 413/18); Hong Kong Epigenomics Project (EpiHK) Fund to H.W. and H.S.; Area of Excellence Scheme (AoE) from RGC (Project number: AoE/M-402/20).

## References

1. J. T. Rodgers et al., mTORC1 controls the adaptive transition of quiescent stem cells from G0 to G(Alert). Nature 510, 393–396 (2014).

2. Y. X. Wang, N. A. Dumont, M. A. Rudnicki, Muscle stem cells at a glance. J Cell Sci 127, 4543–4548 (2014).

3. Y. Li, X. Chen, H. Sun, H. Wang, Long non-coding RNAs in the regulation of skeletal myogenesis and muscle diseases. Cancer Lett 417, 58–64 (2018).

4. Y. Li et al., Long noncoding RNA SAM promotes myoblast proliferation through stabilizing Sugt1 and facilitating kinetochore assembly. Nat Commun 11, 2725 (2020).

5. L. Zhou et al., Linc-YY1 promotes myogenic differentiation and muscle regeneration through an interaction with the transcription factor YY1. Nat Commun 6, 10026 (2015).

6. Y. Zhao et al., MyoD induced enhancer RNA interacts with hnRNPL to activate target gene transcription during myogenic differentiation. Nat Commun 10, 5787 (2019).

7. H. Rahnamoun et al., RNAs interact with BRD4 to promote enhanced chromatin engagement and transcription activation. Nat Struct Mol Biol 25, 687–697 (2018).

8. D. A. Bose et al., RNA Binding to CBP Stimulates Histone Acetylation and Transcription. Cell 168, 135–149 e122 (2017).

9. N. V. N. Carullo et al., Enhancer RNAs predict enhancer-gene regulatory links and are critical for enhancer function in neuronal systems. Nucleic Acids Res 48, 9550–9570 (2020).

10. A. Blank-Giwojna, A. Postepska-Igielska, I. Grummt, lncRNA KHPS1 Activates a Poised Enhancer by Triplex-Dependent Recruitment of Epigenomic Regulators. Cell Rep 26, 2904–2915 e2904 (2019).

11. M. Huarte et al., A large intergenic noncoding RNA induced by p53 mediates global gene repression in the p53 response. Cell 142, 409–419 (2010).

12. N. Dimitrova et al., LincRNA-p21 activates p21 in cis to promote Polycomb target gene expression and to enforce the G1/S checkpoint. Mol Cell 54, 777–790 (2014).

13. E. Hacisuleyman et al., Topological organization of multichromosomal regions by the long intergenic noncoding RNA Firre. Nat Struct Mol Biol 21, 198–206 (2014).

14. J. P. Lewandowski et al., The Firre locus produces a trans-acting RNA molecule that functions in hematopoiesis. Nat Commun 10, 5137 (2019).

15. E. Hacisuleyman, C. J. Shukla, C. L. Weiner, J. L. Rinn, Function and evolution of local repeats in the Firre locus. Nat Commun 7, 11021 (2016).

16. P. F. Tsai et al., A Muscle-Specific Enhancer RNA Mediates Cohesin Recruitment and Regulates Transcription In trans. Mol Cell 71, 129–141 e128 (2018).

17. V. D. Soleimani et al., Transcriptional dominance of Pax7 in adult myogenesis is due to high-affinity recognition of homeodomain motifs. Dev Cell 22, 1208–1220 (2012).

18. N. K. Mullin et al., Wnt/beta-catenin Signaling Pathway Regulates Specific lncRNAs That Impact Dermal Fibroblasts and Skin Fibrosis. Front Genet 8, 183 (2017).

19. L. He et al., In Vivo Study of Key Transcription Factors in Muscle Satellite Cells by CRISPR/Cas9/AAV9-sgRNA Mediated Genome Editing. bioRxiv 10.1101/797746, 797746 (2020).

20. R. W. Yao, Y. Wang, L. L. Chen, Cellular functions of long noncoding RNAs. Nat Cell Biol 21, 542–551 (2019).

21. Y. Huang et al., Large scale RNA-binding proteins/LncRNAs interaction analysis to uncover lncRNA nuclear localization mechanisms. Brief Bioinform 10.1093/bib/bbab195 (2021).

22. D. A. Young et al., Expression of metalloproteinases and inhibitors in the differentiation of P19CL6 cells into cardiac myocytes. Biochem Biophys Res Commun 322, 759–765 (2004).

23. G. Lluri, D. M. Jaworski, Regulation of TIMP-2, MT1-MMP, and MMP-2 expression during C2C12 differentiation. Muscle Nerve 32, 492–499 (2005).

24. A. Bornemann, H. Schmalbruch, Desmin and vimentin in regenerating muscles. Muscle Nerve 15, 14–20 (1992).

25. A. Gallanti et al., Desmin and vimentin as markers of regeneration in muscle diseases. Acta Neuropathol 85, 88–92 (1992).

26. F. Soglia et al., Distribution and Expression of Vimentin and Desmin in Broiler Pectoralis major Affected by the Growth-Related Muscular Abnormalities. Front Physiol 10, 1581 (2019).

27. T. Frohlich et al., Progressive muscle proteome changes in a clinically relevant pig model of Duchenne muscular dystrophy. Sci Rep 6, 33362 (2016).

28. G. Giraud, S. Terrone, C. F. Bourgeois, Functions of DEAD box RNA helicases DDX5 and DDX17 in chromatin organization and transcriptional regulation. BMB Rep 51, 613–622 (2018).

29. H. Yao et al., Mediation of CTCF transcriptional insulation by DEAD-box RNA-binding protein p68 and steroid receptor RNA activator SRA. Genes Dev 24, 2543–2555 (2010).

30. J. Zhou et al., Elevated H3K27ac in aged skeletal muscle leads to increase in extracellular matrix and fibrogenic conversion of muscle satellite cells. Aging Cell 18, e12996 (2019).

31. E. Alessio et al., Single cell analysis reveals the involvement of the long non-coding RNA Pvt1 in the modulation of muscle atrophy and mitochondrial network. Nucleic Acids Res 47, 1653–1670 (2019).

32. W. Zhang, Y. Liu, H. Zhang, Extracellular matrix: an important regulator of cell functions and skeletal muscle development. Cell Biosci 11, 65 (2021).

33. C. W. Guerin, P. C. Holland, Synthesis and secretion of matrix-degrading metalloproteases by human skeletal muscle satellite cells. Dev Dyn 202, 91–99 (1995).

34. G. Fibbi et al., Cell invasion is affected by differential expression of the urokinase plasminogen activator/urokinase plasminogen activator receptor system in muscle satellite cells from normal and dystrophic patients. Lab Invest 81, 27–39 (2001).

35. G. Pallafacchina et al., An adult tissue-specific stem cell in its niche: a gene profiling analysis of in vivo quiescent and activated muscle satellite cells. Stem Cell Res 4, 77–91 (2010).

36. F. Relaix et al., Perspectives on skeletal muscle stem cells. Nat Commun 12, 692 (2021).

37. H. Yamakawa, D. Kusumoto, H. Hashimoto, S. Yuasa, Stem Cell Aging in Skeletal Muscle Regeneration and Disease. Int J Mol Sci 21 (2020).

38. Y. Zheng, T. Liu, Q. Li, J. Li, Integrated analysis of long non-coding RNAs (lncRNAs) and mRNA expression profiles identifies lncRNA PRKG1-AS1 playing important roles in skeletal muscle aging. Aging (Albany NY) 13, 15044–15060 (2021).

39. V. Sartorelli, S. M. Lauberth, Enhancer RNAs are an important regulatory layer of the epigenome. Nat Struct Mol Biol 27, 521–528 (2020).

40. C. L. Hsieh et al., Enhancer RNAs participate in androgen receptor-driven looping that selectively enhances gene activation. Proc Natl Acad Sci U S A 111, 7319–7324 (2014).

41. G. Arun, V. S. Akhade, S. Donakonda, M. R. Rao, mrhl RNA, a long noncoding RNA, negatively regulates Wnt signaling through its protein partner Ddx5/p68 in mouse spermatogonial cells. Mol Cell Biol 32, 3140–3152 (2012).

42. G. Caretti et al., The RNA helicases p68/p72 and the noncoding RNA SRA are coregulators of MyoD and skeletal muscle differentiation. Dev Cell 11, 547–560 (2006).

43. R. Sambasivan et al., Distinct regulatory cascades govern extraocular and pharyngeal arch muscle progenitor cell fates. Dev Cell 16, 810–821 (2009).

44. F. Chen et al., YY1 regulates skeletal muscle regeneration through controlling metabolic reprogramming of satellite cells. The EMBO journal 38 (2019).

45. X. Chen et al., Malat1 regulates myogenic differentiation and muscle regeneration through modulating MyoD transcriptional activity. Cell Discov 3, 17002 (2017).

46. X. L. Peng et al., MyoD- and FoxO3-mediated hotspot interaction orchestrates super-enhancer activity during myogenic differentiation. Nucleic Acids Res 45, 8785–8805 (2017).

47. K. K. So, X. L. Peng, H. Sun, H. Wang, Whole Genome Chromatin IP-Sequencing (ChIP-Seq) in Skeletal Muscle Cells. Methods Mol Biol 1668, 15–25 (2017).

48. B. Langmead, S. L. Salzberg, Fast gapped-read alignment with Bowtie 2. Nat Methods 9, 357–359 (2012).

49. Y. Zhang et al., Model-based analysis of ChIP-Seq (MACS). Genome Biol 9, R137 (2008).

50. R. L. Williams, Jr. et al., fourSig: a method for determining chromosomal interactions in 4C-Seq data. Nucleic Acids Res 42, e68 (2014).

