## Supplementary information for "seRNA *PAM-1* regulates skeletal muscle satellite cell activation and aging through *trans* regulation of *Timp2* expression synergistically with Ddx5"

**Inventory of Supplementary Information**

**1. Supplementary Figures**

Figure S1. lncRNAs profiling identifies *PAM-1* as a seRNA promoting SC activation.

Figure S2. *PAM-1* interacts with inter-chromosomal loci to modulate target expression

**2. Supplementary Tables**

Table S1. List of differentially expressed lncRNAs during SC activation.

Table S2. List of *PAM-1* seRNA targets in ChIRP-seq.

Table S3. List of *PAM-1* SE targets in 4C-seq.

Table S4. Oligos used in the study.

**3. Supplementary figure legends**

**Supplementary Figure S1. lncRNAs profiling identifies *PAM-1* as a seRNA promoting SC activation.** **A.** Differentially expressed genes (DEG) analysis of ASC-24h vs. FISC. Yellow dots indicate differentially expressed lncRNAs. *PAM-1* is highly expressed in ASC versus FISC. **B.** List of top ranked lncRNAs according to fold change in expression level between ASC-24h and FISC. **C.** Examples of top ranked differentially expressed lncRNAs.

**Supplementary Figure S2**. ***PAM-1* interacts with inter-chromosomal loci to modulate target expression**. **A.** Individual Gene Ontology (GO) terms of genome-wide common targets of *PAM-1* identified in Figure 2I. **B.** Genome browser tracks showing H3K27ac ChIP-seq peaks along *PAM-1* locus. Design and selection of sgRNAs targeting *PAM-1* locus is shown. sgRNA1 and sgRNA2 were designed to delete a 59 bp region on *PAM-1*.
