## Supplementary Figures for "seRNA *PAM-1* regulates skeletal muscle satellite cell activation and aging through *trans* regulation of *Timp2* expression synergistically with Ddx5"

**Supplementary Figure S1. So K.K.H. & Huang Y. et. al.**

**A**

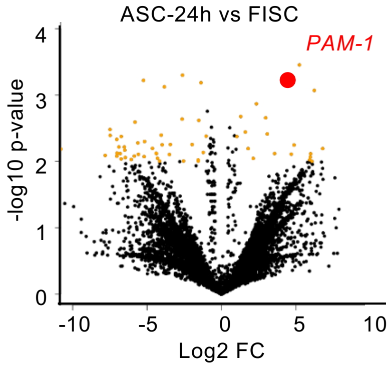


**B**

| Ref Seq ID | FISC | ASC-24h |
|---|---|---|
| Gm12603 (*PAM-1*) | 0 | 12.719 |
| 4632415L05Rik | 0.420 | 3.731 |
| 1810058I24Rik | 3.617 | 12.521 |
| CUFF.5420 | 0.101 | 1.812 |
| CUFF.5390 | 0.036 | 1.547 |

**C**

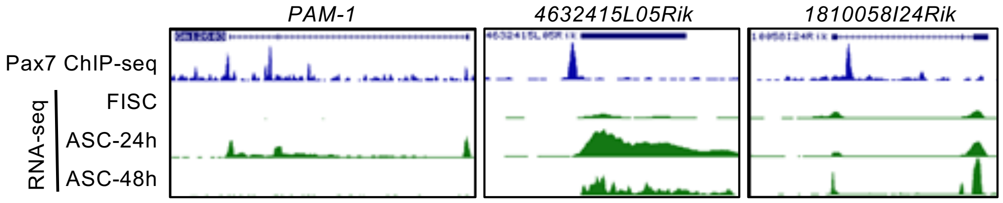

### Supplementary Figure S2. So K.K.H. & Huang Y. et. al.

A

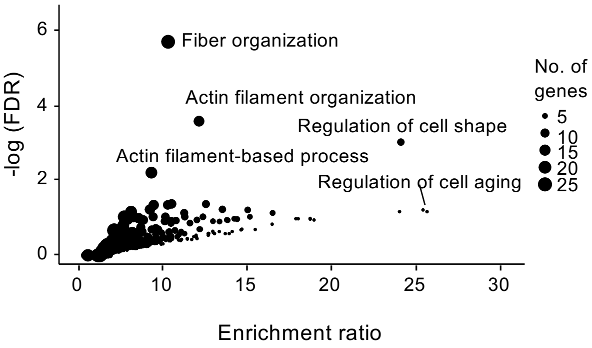


B

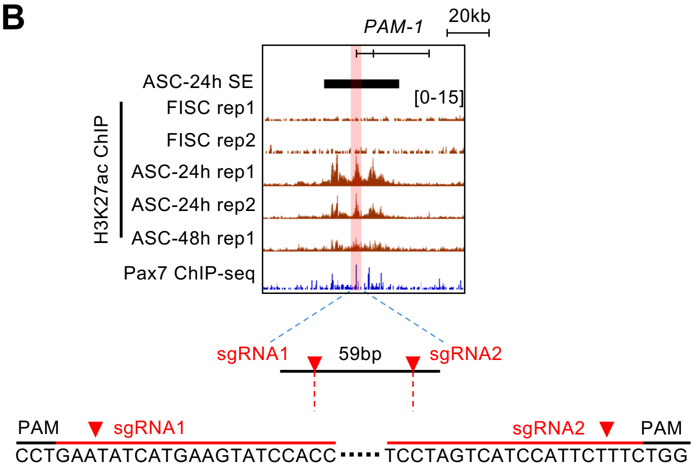
